# Phosphoproteomics reveals novel BCR::ABL1-independent mechanisms of resistance in chronic myeloid leukemia

**DOI:** 10.1101/2025.01.08.631869

**Authors:** Valeria Bica, Veronica Venafra, Giorgia Massacci, Simone Graziosi, Sara Gualdi, Gessica Minnella, Federica Sorà, Patrizia Chiusolo, Maria Elsa Brunetti, Gennaro Napolitano, Massimo Breccia, Dimitrios Mougiakakos, Martin Böettcher, Thomas Fischer, Livia Perfetto, Francesca Sacco

**Affiliations:** Ph.D. Program in Cellular and Molecular Biology, Department of Biology, University of Rome ‘Tor Vergata’, Rome, Italy; Department of Biology, University of Rome Tor Vergata, Rome, Italy; Department of Biology and Biotechnologies “C. Darwin”, University of Rome La Sapienza, Rome, Italy; Sezione di Ematologia, Dipartimento di Scienze Radiologiche ed Ematologiche, Università Cattolica del Sacro Cuore, Roma, Italy; Dipartimento di Scienze di laboratorio ed Ematologiche, Fondazione Policlinico A. Gemelli IRCCS, Roma; Telethon Institute of Genetics and Medicine (TIGEM), Naples, Italy; Department of Medical and Translational Science, Federico II University, Naples, Italy; Scuola Superiore Meridionale (SSM), School for Advanced Studies, Genomics and Experimental Medicine program, Naples, Italy; Department of Translational and Precision Medicine, Azienda Policlinico Umberto I, Sapienza University of Rome, Rome, Italy; Health campus for Inflammation, Immunity and Infection (GCI3), University of Magdeburg, Germany; Department of Hematology, Oncology and Cell Therapy, University of Magdeburg, Magdeburg, Germany

## Abstract

BCR::ABL1 drives chronic myeloid leukemia (CML) disease and treatment, as revealed by the success of tyrosine kinase inhibitor (TKI) therapy. However, additional poorly characterized molecular pathways, acting as BCR::ABL1 independent mechanisms, play crucial roles in CML, contributing to leukemic stem cells (LSCs) persistence, TKI resistance and disease progression. Here, by combining high sensitive mass spectrometry (MS)-based phosphoproteomics with the *SignalingProfiler* pipeline, we obtained two signaling maps offering a comprehensive description of the BCR::ABL1 dependent and independent pro-survival signalling mechanisms. We leveraged these maps to unbiasedly and systematically discover therapeutic vulnerabilities, by implementing the *Druggability Score* computational algorithm. By this strategy, and in combination with *in vitro* and in *ex vivo* functional assays, we show a crucial role of acquired FLT3-dependency in resistant CML models. In conclusion, we reposition FLT3, one of the most frequently mutated drivers of acute leukemia, as a potential therapeutic target for TKI resistant CML patients.

## Introduction

Chronic Myeloid Leukemia (CML) is a clonal myeloproliferative disorder, molecularly defined by the presence of the Philadelphia chromosome (Ph), resulting from a reciprocal translocation between chromosomes 9 and 22 [t(9;22)(q34;q11)]^1^. This translocation leads to the formation of the BCR::ABL1 chimeric protein, a constitutively active tyrosine kinase that drives aberrant proliferation and survival, promoting leukemogenesis^2^. Given the pivotal role of BCR::ABL1 in CML pathogenesis, tyrosine kinase inhibitors (TKIs), such as imatinib, were developed, markedly improving patient survival^3,4^. However, TKI therapy fails in about 30% of newly diagnosed CML patients, due to different mechanisms of resistance^5^. TKI resistance mechanisms can be broadly classified as BCR::ABL1-dependent or BCR::ABL1-independent^6^. CML non-responder patients with BCR::ABL1-dependent resistance are typically characterized by the acquisition of point mutations in the kinase domain or overexpression of the oncogene^7^ and can receive specific TKIs according to mutational assessment. In contrast, 50% or more of non-responder patients do not harbour BCR::ABL1 mutations and lack therapeutic strategies, as the basis of such BCR::ABL1–independent resistance remains poorly understood^8–10^. BCR::ABL1 independent resistance mechanisms also play a crucial role in leukemia stem cells (LSCs), which are intrinsically resistant to TKI therapy and hinder long-term discontinuation, namely treatment-free-remission, in responder patients^11,12^. In our study, we aim to elucidate the BCR::ABL1 dependent and independent resistance mechanisms, with the ultimate goal of proposing novel therapeutic strategies to counteract resistance and improve clinical outcomes of relapsed or unresponsive CML patients. To achieve this, we employed an integrated, system-based strategy combining high-resolution mass spectrometry-based (phospho)proteomics of imatinib responsive and unresponsive cell lines with transcriptomic analysis of primary blasts derived from responsive and unresponsive CML patients, and a network-based approach. By this strategy, we derived two comprehensive BCR::ABL1-dependent and BCR::ABL1-independent signaling maps, identifying novel players in imatinib response and deciphering the complex signaling pathways rewiring occurring in CML resistant cells. By implementing the *Druggability Score* algorithm, we leveraged these networks to identify and validate promising drug targets killing resistant cells. *Ex vivo* validation in LSCs highlighted a crucial role of acquired FLT3-dependency in resistant CML models. Finally, our study provides insights into the mechanisms underlying imatinib response and resistance and identifies FLT3-TKIs e.g. midostaurin as a potential effective therapeutic strategy for unresponsive patients.

## Results

### Multi-omic analysis of imatinib sensitive and resistant CML cells

To characterize the BCR::ABL1 dependent and independent signaling pathways promoting cell survival, we developed a CML model of imatinib-resistant cells (**Fig.1A**). Specifically, we established a K562 cell line selected to persist upon chronic imatinib exposure (K562-R) (**Fig.S1A**). The persistence of these cells is accompanied by a drastic reduction of the BCR::ABL1 expression level and its downstream proteins (**Fig.S1B-C**), without any impact on their cell cycle progression (**Fig.S1D**), implying that BCR::ABL1 independent signaling pathways may promote cell survival. This celluar model, together with imatinib-sensitive CML cell lines (K562 and LAMA84), was profiled by state-of-the-art MS-based (phospho)proteomics after the incubation with imatinib for 24h (**Table S1** and **S2**). By this strategy, more than 7,000 proteins and 19,000 class I phosphosites were quantified in biological quadruplicates (**Fig.1B,S2A**). Next, to assess how the (phospho)proteome was changed in imatinib treated and resistant cells, we extracted significantly deregulated proteins and phosphosites by t-test analysis (FDR < 0.05), comparing: i) K562-Ima vs K562 (control cells), ii) LAMA84-Ima vs LAMA84; and iii) K562-R vs K562. Imatinib treatment remodels approximately 30% of the phosphosites and proteins in K562 and LAMA84 cell lines, while more than 70% of the phosphoproteome and the proteome (15,000 phosphosites and 5,000 proteins) is significantly modulated in K562-R cells as compared to control cells (**Fig.S2B**). To note, the impact of imatinib on the (phospho)proteome is highly reproducible in K562 and LAMA84 cell lines (R = 0.7) (**Fig.S2C**). In addition, imatinib induces a consistent modulation of phosphorylation and protein abundance in 20% of the phosphosites (**Fig.S2B**).

**Figure 1.**
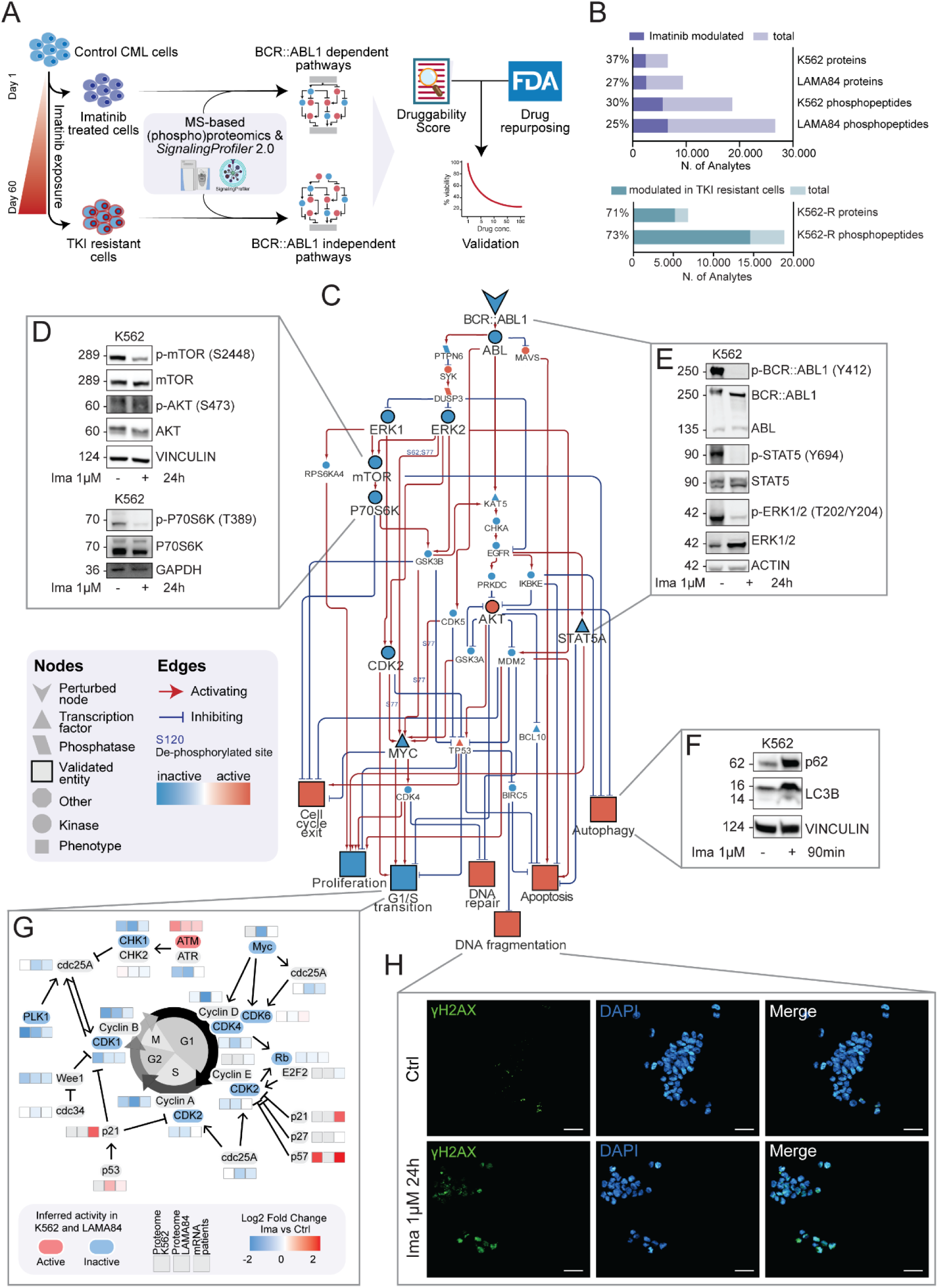
Characterization of BCR::ABL1 dependent mechanisms in sensitive cells. **A.** Imatinib sensitive and resistant cells underwent MS-based (phospho)proteomics. *SignalingProfiler* 2.0 was employed to derive context-specific signaling networks describing BCR::ABL1 dependent and independent signaling pathways. The *Druggability Score* pipeline was employed to repurpose FDA-approved drugs triggering resistant cells’ death. In vitro and ex vivo validation was performed to identify novel therapeutic strategies for unresponsive CML patients. **B.** Quantification coverage of phosphopeptides and proteins. For K562 and LAMA84 cells perturbed by 24h of imatinib 1µM treatment, significantly modulated analytes (t test, S0=0.1, FDR<0.05) are reported in violet (upper panel). For K562-R cells compared to K562 cells, significantly modulated analytes (t test, S0=0.1, FDR<0.05) are reported in green (lower panel). **C.** Functional submodel extracted from *SignalingProfiler* 2.0 output linking BCR::ABL1 to cellular phenotypes modulated upon imatinib treatment. **D.** Representative western blots of PI3K/AKT/mTOR axis activity upon 24h of 1µM imatinib treatment. **E.** Representative western blots of BCR::ABL1 activity status, MAPK and JAK/STAT canonical downstream pathways upon 24h of 1µM imatinib treatment. **F.** Representative western blots showing expression levels of key autophagy regulators, such as p62 and LC3B perturbed by 90 minutes of imatinib 1µM treatment. **G.** Illustration showing cell cycle-related proteins modulation by imatinib at mRNA, protein, and activation level. **H.** Representative images of γH2AX staining in K562 cells perturbed with imatinib for 24 hours. Scale bars represent 50 μm.

To assess the clinical relevance of our experimental models, we took advantage of two recently published transcriptome datasets comparing i) the gene expression modulation of imatinib-treated patients to untreated ones (GSE216837); and ii) non-responder patients to the responders’ counterpart (GSE14671) (**Fig.S2D, Table S1, S2**). Thus, we investigated how the transcriptome, proteome and phosphoproteome rewiring in imatinib treated and resistant CML models impacted crucial biological processes by enrichment analysis. Briefly, proteins involved in metabolic processes, such as TCA, glycolysis/gluconeogenesis are significantly over-expressed and hyperphosphorylated in both imatinib treated and resistant cells (**Fig.S2E**). Consistently with its involvement in cancer therapeutic resistance, the arachidonic metabolism is significantly upregulated in both K562-R cells and non-responder patients at the protein and mRNA level, respectively^13^. Finally, in line with the pro-apoptotic effect of imatinib, pro-survival, and proliferation signaling pathways, such as JAK/STAT, MAPK, and mTOR, are significantly downregulated through phosphorylation-based mechanisms only in imatinib treated K562 and LAMA84 cells. Consistently, cell cycle-related proteins are down-regulated only in imatinib-treated CML models. Taken together, these observations indicate that the imatinib-induced response in LAMA84 and K562 is consistent and robust, and confirm the relevance of our cell lines as models to elucidate the BCR::ABL1 dependent and independent proliferative pathways.

### Identifying BCR::ABL1-dependent pro-survival signaling pathways

First, we aimed to systematically describe BCR::ABL1 dependent survival pathways suppressed by imatinib treatment. To this aim, by employing *SignalingProfiler* 2.0^14^, we inferred the activity of signaling proteins in imatinib-treated K562 and LAMA84 cells as compared to control cells. As expected, the treatment similarly impacts the activity of kinases, phosphatases, and transcription factors in K562 and LAMA84 cells (R=0.48, p=2.2·10^-16^) (**Fig.S3A**). In particular, imatinib inactivates ABL1 and its well-characterized downstream targets (STAT5A, RPS6) as well as cyclin-dependent kinases, such as CDK2, CDK4, CDK5, and CDK6. Our approach also enables the identification of kinases whose activity is increased in this condition. Many of them are involved in the DNA damage pathway, including ATM and members of the NEK family kinase (e.g. NEK2, NEK5, NEK9). Importantly, by applying *SignalingProfiler* 2.0 to the patient transcriptome (imatinib-treated cohort), we found that the activity of most of the identified transcription factors, including the cell-cycle regulator MYC, is similarly regulated in CML patients and cell lines (R=0.78, p=0.039) (**Fig.S3B-C**), thus reassuring about the translational impact of the approach. Next, we considered proteins with consistent activity modulation upon imatinib treatment in K562 and LAMA84 cells (50 transcription factors, 81 kinases and phosphatases, and 451 other signaling proteins - **Table S3**), and generated the BCR::ABL1 dependent-signaling network. The resulting graph accounts for 200 nodes and 524 edges, suggesting a deep signaling rewiring induced by the drug. Interestingly, the phenotypic inference indicates that imatinib activates apoptosis, DNA fragmentation, autophagy and repair, and inhibits cell cycle progression (e.g., G1/S transition) in both cell lines (**Fig.S3D**). To reduce the complexity of this map, we selected the functional circuits linking BCR::ABL1 to the predicted phenotypes (**Fig.1C**). Remarkably, the network-based approach and the *in vitro* validation experiments align closely and recapitulate the well-characterized relations between BCR::ABL1 and its downstream signaling pathways (ERK1/2, STAT5 and mTOR) and cell-cycle regulation (**Fig.1D-E**). Additionally, the map generates novel mechanistic hypotheses about the signaling events perturbed by imatinib exposure. Interestingly, while mTOR and its downstream targets, such as p70S6K, are dephosphorylated and inhibited upon imatinib treatment, feedback loops hyperactivate AKT (**Fig.1D**). In line with mTOR pathway inhibition, LC3-II and p62 abundance confirmed an increase in autophagy induction (**Fig.1F**). The inspection of the imatinib-dependent modulation of cell cycle reveals that crucial proteins, including CDKs, Rb and ATM kinase are significantly and consistently modulated at multiple regulatory layers in cell lines and patients (**Fig.1G**) driving G1 phase arrest with consequent inhibition of proliferation, as confirmed by functional assays (**Fig.S1D** and **S3E**). In line with the *in silico* activation of DNA fragmentation and repairs phenotypes (**Fig.1C** and **S3D**), we detected higher γH2AX levels in imatinib-treated cells (**Fig.1H**). Altogether our data indicate that imatinib-dependent suppression of BCR::ABL1 triggers apoptosis by globally rewiring signaling pathways through DNA damage induction and cell cycle block.

### Identifying BCR::ABL1 independent pro-survival signaling pathways

Here we aim to obtain a mechanistic picture of the BCR::ABL1-independent signaling remodeling. Thus, we employed our cell line models, namely K562-R and K562-Ima cells, wherein BCR::ABL1 is suppressed, through transcriptional and pharmacological mechanisms, respectively. Interestingly, the two models display divergent behaviour as the first promotes cell survival, whereas the second triggers cell death. This offers the unprecedented opportunity to discover commonly regulated signaling axes, as well as compensative and divergent resistance mechanisms. First, we ran *SignalingProfiler* 2.0 to derive the activity of 261 transcription factors, 280 kinases and phosphatases, and 2153 other signaling proteins in K562-R as compared to control cells (**Table S4**). As expected, we observe a poor correlation between the inferred activity in resistant cells with respect to imatinib-treated cells (**Fig.2A-B**). Most of the proteins (2470/3005 proteins), including PLK1, LYN, and MYC, are oppositely regulated, supporting the hypothesis that proliferation and survival of K562-R cells rely on BCR::ABL1 independent mechanisms. Next, we employed *SignalingProfiler* 2.0 to derive the K562-R specific network. This resulted in a graph with 700 proteins and 10 phenotypes linked by 1985 interactions, with 20% of phosphorylation events experimentally quantified (**Table S4**), revealing a huge reorganization in BCR::ABL1-depleted resistant cells. As quality control, we compared our results with genes previously implicated in imatinib resistance by an independent genome-wide CRISPR-Cas9 screening^15^. Remarkably, 70% of the hits are consistently down-regulated and connected in the network (**Fig.S4A**), supporting the ability of our network-based approach to identify drug resistance pathways. Next, we compared the BCR::ABL1 dependent network (**Fig.1C**) with the newly generated K562-R specific map. We observe that BCR::ABL1 downstream signaling effectors, namely STAT5A and CDK1/4, as well as the DNA damage response are equally modulated in both models (**Fig.S4B-C**), suggesting the persistence of mechanisms associated with the inhibition of BCR::ABL1. To identify BCR::ABL1 independent pro-proliferative molecular mechanisms, we extracted the subnetwork downstream of receptors oppositely modulated in K562-R and imatinib-treated K562 cells (**Fig.S4D),** and impacting on proliferation and apoptosis (**Fig.2C, Table S5**). The BCR::ABL1 independent-specific subnetwork as well as validation assays indicate a complex rewiring of the mTOR pathway (**Fig.2C-D**). While mTOR is down-regulated, the activity of its canonical targets (e.g. P70S6K) as well as the abundance of the upstream PI3K kinase (p85α, regulatory subunit) is increased in K562-R as compared to control cells (**Fig.2D**). As revealed by increased activity of BCL2 and inhibition of TP53 and BID, the apoptotic pathway is suppressed. Also, transcription factors MYC and RELA appear consistently regulated by *in silico* prediction and experimental validation (**Fig.2G-H**). Interestingly, both oncogenic kinases JAK1 and FLT3 are up-regulated at protein and phosphorylation levels (**Fig.2E-I**). Indeed, both FLT3 and JAK1/2 are frequently mutated in haematological disorders and targeted by drugs currently approved in clinics^16^. Taken together, these observations suggest that several signaling proteins, including JAK1 and FLT3, promote cell proliferation, in absence of BCR::ABL1 (**Fig.S1A**), representing promising drug targets.

**Figure 2.**
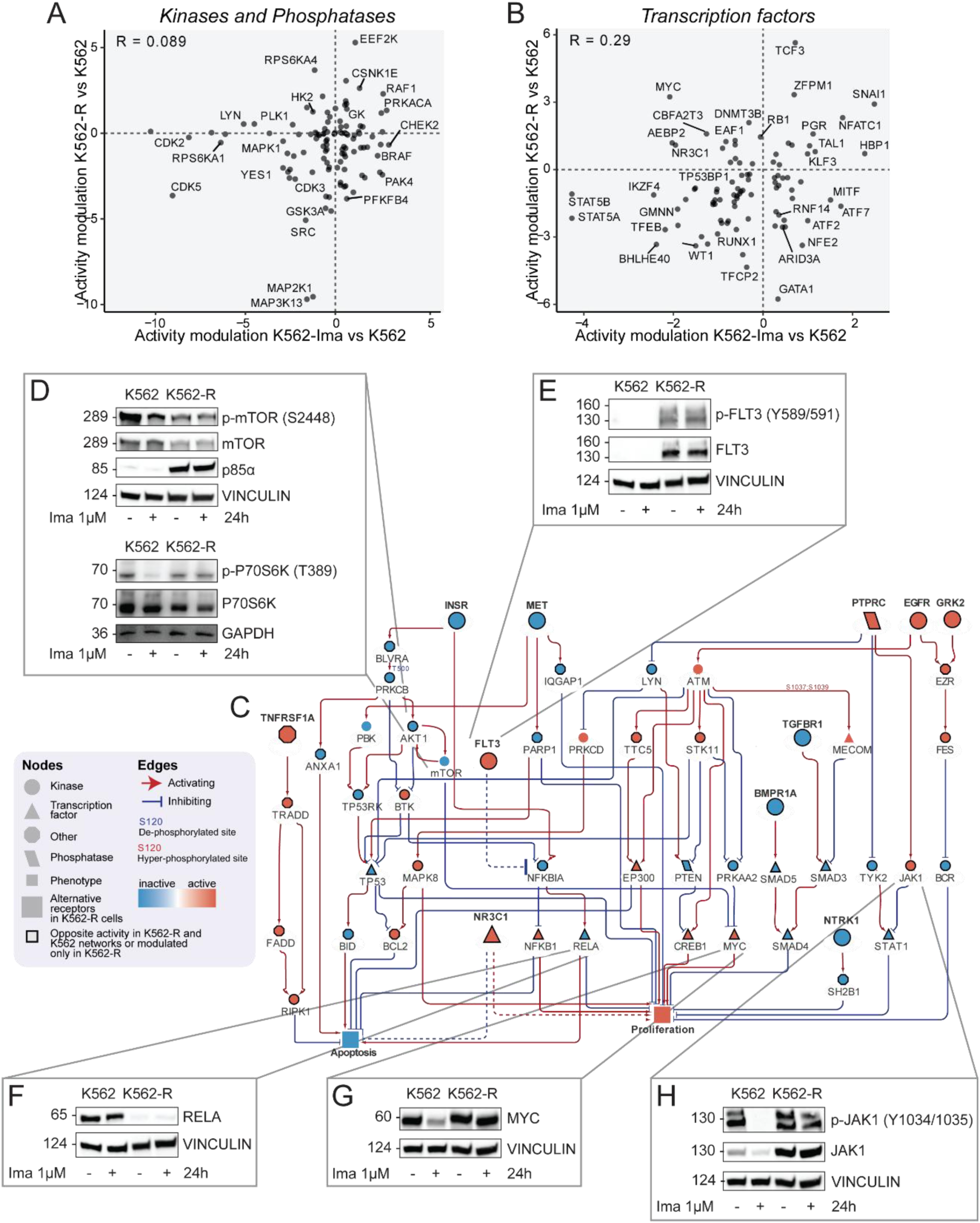
Characterization of BCR::ABL1 independent mechanisms in resistant cells. **A-B.** Scatterplots showing the comparison between kinases/phosphatases (**A**) and transcription factors (**B**) activity predicted in K562 and K562-R cells vs control cells datasets for kinases and phosphatases. R indicates Pearson correlation. **C.** Functional submodel extracted from BCR::ABL1 independent-specific subnetwork reporting paths going from 10 alternative receptors oppositely modulated in imatinib K562-R and K562 cells exposed to imatinib to apoptosis and proliferation phenotypes. **D.** Representative western blots of PI3K/AKT/mTOR axis activity upon 24h of 1µM imatinib treatment in control cells and K562-R cells. **E-F.** Representative western blot of control cells and K562-R cells showing phosphorylation status and protein abundance of FLT3 receptor (**E**) and RELA protein (**F**) upon imatinib treatment. Both panels were obtained from the same gel and divided for aesthetic purposes of the figure. **G-H.** Representative western blot of control cells and K562-R cells showing phosphorylation status and protein abundance of MYC transcription factor (**G**) and JAK1 kinase (**H**) upon imatinib treatment. Both panels were obtained from the same gel and divided for aesthetic purposes of the figure.

### Repurposing FDA drugs to eradicate resistant CML cells

To unbiasedly identify druggable targets killing K562-R cells, we implemented the *Druggability Score* algorithm (**Fig.3A**), ranking each of the nodes in the BCR::ABL1 independent network according to the following criteria: (i) activation level in K562-R cells; (ii) number of paths impacting on apoptosis and proliferation; (iii) degree centrality (**Table S6**). Proteins with a positive *Druggability Score* are expected to induce greater cell death in resistant cells compared to control cells and *vice versa*. We focused on the eight proteins with the highest *Druggability Score* (FLT3, JAK1, BTK, PIK3CB, PIK3C3, DNMT3A, DNMT1, and BCL2) targeted by FDA-approved drugs for hematological malignancies^16^ (**Fig.3B**). Additionally, as negative control, we also included AKT, whose *Druggability Score* is negative. FDA-approved inhibitors of these eight proteins were used to treat resistant and sensitive K562 cells and cell viability was assessed by MTT assay (**Fig.3C**). Our *in vitro* results closely align with the prediction: drugs with a positive *Druggability Score* induce greater cell death in resistant cells as compared to sensitive cells. As expected, pharmacological inhibition of AKT leads to higher cell death in sensitive cells than in resistant ones. Strikingly, FLT3 inhibition by two different drugs (quizartinib and midostaurin) emerged as the most effective strategy for eradicating resistant CML cells, as also demonstrated by their IC50 values (**Fig.3C** and **S4E**). As previously described, leukemic stem cells (LSCs) are intrinsically resistant to BCR::ABL1 inhibition^17^. Thus, we asked whether LSCs, isolated from three CML patients, could benefit from pharmacological suppression of FLT3. LSCs and leukemic progenitor cells (LPCs) were FACS-sorted (**Fig.S6**) and treated with midostaurin for 24h, followed by cell viability assessment. Indeed, pharmacological suppression of FLT3 more effectively killed LSCs than LPCs (**Fig.3D**, **Table S7**). In agreement with these results, we found that FLT3 was not only upregulated in K562-R cells at the protein and phosphorylation levels (**Fig.2E**), but also at the mRNA level in LSCs as compared to LPCs, as revealed in an independent transcriptome dataset of a cohort of CML patients (GSE43754) (**Fig.3E**). Prompted by these findings, we also investigated whether FLT3 expression upregulation could also be detected in non-responder patients. Although not significant, the expression of FLT3 is higher in non-responder patients as compared to responders’ counterparts (**Fig.S4F**). This is consistent also with another CML patient dataset from the GEO database (**Fig.S4G**). In conclusion, FLT3 inhibition emerges as an effective strategy to overcome BCR::ABL1 independent imatinib resistance in CML.

**Figure 3.**
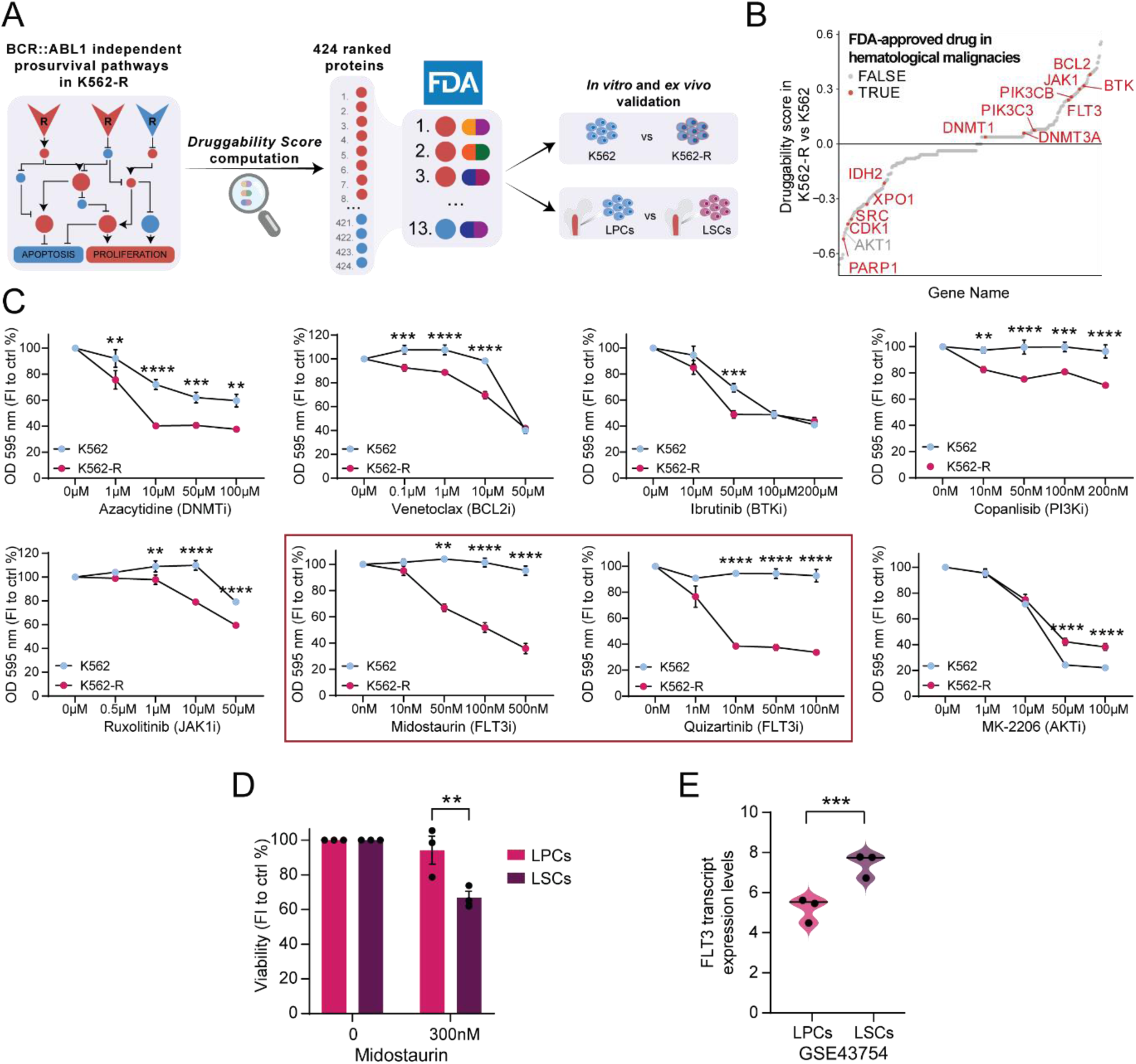
Identification of new druggable targets and repurposing of FDA-approved drugs. **A.** Druggable targets prioritization strategy. **B.** Scatterplot showing drug targets ranked according to the *Druggability Score*. **C.** MTT assay on K562 and K562-R cells exposed for 24h at different concentrations of FDA-approved inhibitors of prioritized targets. MK-2206 (AKTi) was used as negative control. The graphs show the percentage of absorbance at 595 nm normalized on control condition. The reported statistical significance is between K562 and K562-R cells at the same experimental condition. **D.** LPCs (CD38+) and LSCs (CD26+) were sorted and exposed to midostaurin for 24h. Viability assay was performed by trypan blue exclusion. **E.** Quantification of FLT3 transcript levels in LPCs and LSCs from RNAseq analysis obtained from GEO dataset (GSE43754).

## Discussion

Imatinib, known as the “magic bullet”, thanks to its ability to shut-down BCR::ABL1 activity, has revolutionized the treatment landscape of CML. However, tyrosine-kinase inhibitor (TKI) therapy fails in about 30% of newly diagnosed CML patients, which develop different mechanisms of therapy resistance^5^. Additionally, approximately 50% of CML responder patients achieve treatment-free remission due to difficulty in completely eradicating LSCs, which are intrinsically resistant to TKIs and are the reservoir for disease persistence^18,19^. These observations pose complex therapeutic questions: for TKI responders, how can we completely eradicate leukemia stem cells? For pan TKI non-responders, what are alternative therapies that can be employed to treat CML? To address these questions, we performed a large-scale integrated analysis, combining state-of-the-art mass spectrometry-based phosphoproteomics of imatinib responsive and unresponsive cell lines with transcriptomic studies of responsive and unresponsive patient-derived primary blasts and a network-based approach. Hence, we obtained a comprehensive description of the BCR::ABL1 dependent and independent pro-survival signaling mechanisms. With more than 25,000 phosphosites and 8,000 proteins accurately quantified in imatinib treated and resistant cell lines, our large-scale multi-layered dataset represents an unprecedented resource for the scientific community^20,21^. By employing our in-house implemented *SignalingProfiler* 2.0 pipeline, we derived the molecular paths through which BCR::ABL1 controls crucial phenotypes including cell cycle, autophagy, protein synthesis, and DNA damage^22–25^. Remarkably, our strategy offered an in-depth description of imatinib-dependent modulation of cell cycle, as most of its regulators were coherently measured at proteome, transcriptome, and activity levels, in cell lines and in patient-derived primary blasts. Next, we aimed at narrowing down alternative therapeutic strategies overcoming TKI resistance and eradicating LSCs. The comparison between resistant and control cells revealed a huge remodeling of crucial processes at both the proteome and phosphoproteome regulatory layers. We integrated our data with *SignalingProfiler* 2.0 and derived the BCR::ABL1 independent signaling network. Interestingly, crucial membrane receptor kinases, including FLT3, EGFR, GRK2 and TNFR are hyperactive in the newly generated resistant network and are potentially implicated in sustaining pro-proliferative pathways in absence of BCR::ABL1. Thus, the generated maps not only provide a comprehensive description of the molecular mechanisms implicated in TKI-resistance, but can be leveraged to unbiasedly and systematically rank druggable targets. Indeed, we implemented the *Druggability Score*, a generally applicable algorithm which repurposes FDA approved drugs, by prioritizing nodes, according to their topological properties and activation levels in resistant models. This strategy allowed us to pinpoint and *in vitro* validate BCL2, JAK1, BTK, FLT3, PI3KCB, PI3KC3, DNMT1, and DNMT3A as candidate targets. Interestingly, a recent study also identified BCL2 and JAK1 as key proteins involved in TKI resistance^26^, suggesting the reliability of our strategy. In our analysis, the FLT3 inhibition by two commonly used FDA approved drugs (midostaurin and quizartinib) emerged as the most effective solution to eradicate resistant cells. FLT3 is a tyrosine-kinase receptor frequently mutated in hematological disorders and associated with dismal prognosis^27^. Remarkably, we and others observed that FLT3 is upregulated at multiple levels in different TKI resistance models of CML: i) protein abundance and activity in K562-R cell line; ii) transcript level in blasts derived from an independent small cohort of TKI non-responders patients profiled in this study; iii) transcript level in blasts derived from a large cohort of blast-phase patients^28^; and iv) transcript level in patient-derived leukemic stem cells (LSCs)^29^. These observations prompted us to investigate the effect of midostaurin treatment on LSCs. Remarkably, pharmacological suppression of FLT3 effectively killed LSCs as compared to leukemic progenitors. Altogether, these results clearly indicate a crucial role of acquired FLT3-dependency in resistant CML models. Mechanistically, FLT3 has been implicated in TKI resistance through the JAK-STAT axis^30^. However, we and others show that JAK suppression has a mild impact on triggering cell death in K562-R and in patient-derived LSCs^26^, suggesting alternative FLT3-mediated axes. Our map suggests the potential implication of the NFkB transcription factor in FLT3-dependent TKI resistance. In conclusion, the combination of MS-based (phospho)proteomics and network-based approaches enabled repositioning FLT3, one of the most frequently mutated drivers of leukemia, as a therapeutic target for unresponsive patients and prognostic markers of TKI resistance.

## Data availability

The mass spectrometry proteomics and phosphoproteomics data have been deposited to the ProteomeXchange Consortium via the PRIDE^31^ partner repository with the dataset identifier PXD056957. Patients’ transcriptomic data are available on the GEO database under the ID: GSE280476. All code used for the network generation and *Druggability Score* analysis is available at https://github.com/SaccoPerfettoLab/Chronic_myeloid_leukemia_SignalingProfiler2.0_analysis.

## Acknowledgments

We thank Dr. Serena Paoluzi and Dr. Marta Iannuccelli for their technical support. This research was funded by the Italian Association for Cancer Research (AIRC) with a grant to L.P. (MFAG Grant n. 28858) and a grant to F.S. (Start-Up Grant n. 21815) and by MUR PRIN 2022 (n. E53D23004850006). G.M. is supported by MUR PRIN 2022 (n. E53D23004850006). L.P. and F.S. are supported by a joint PRIN 2022 PNRR grant (n. P2022JRETW), funded by the European Union – NextGenerationEU, and by a SEED Sapienza Grant. V.V is supported by PON-MUR fellowship (n. DOT13IEP1U-1)

## Author contributions

Conceptualization, V.B., V.V., L.P., F.S.; methodology, V.B., V.V., G.M., S.G., L.P., F.S.; formal analysis, V.B., V.V., G.M., S.G., P.C., M.B., D.M.; investigation, V.B., V.V., with the contribution of T.F., M.B., D.M.; writing original draft preparation, V.B., V.V., L.P., F.S.; writing review and editing, all; supervision, L.P., F.S.; funding acquisition, F.S. All authors have read and agreed to the published version of the manuscript.

## Declaration of interest

G.N. is advisory board member of Amplify Therapeutics.

## Supplemental information

**Document S1.** Figures S1-S6

**Table S1.** Excel file containing proteomics and phosphoproteomics profile of K562 and LAMA84 cell lines and transcriptomics profile of Chronic Myeloid Leukemia patients from GEO database (ID: GSE216837) upon 24 imatinib treatment.

**Table S2.** Excel file containing baseline proteomics and phosphoproteomics profiles of K562-R and K562 cells and transcriptomics profile of Chronic Myeloid Leukemia patients from GEO database (ID: GSE14671).

**Table S3.** Excel file containing the list of inferred proteins, nodes and edges of the signaling network generated by *SignalingProfiler* 2.0 from the (phospho)proteomics profile of K562 and LAMA64 cell lines upon 24h imatinib treatment.

**Table S4.** Excel file containing the list of inferred proteins, nodes and edges of the signaling network generated by *SignalingProfiler* 2.0 from the baseline (phospho)proteomics profile of K562-R and K562 cell lines.

**Table S5.** Excel file containing nodes and edges of the functional circuit extracted from the BCR::ABL1 independent signaling network connecting 23 oppositely modulated receptors to cancer hallmarks phenotypes.

**Table S6.** Excel file containing BCR::ABL1 independent signaling subnetwork nodes ranked according to the topological properties of the Druggability Score algorithm.

**Table S7.** Excel file containing clinical characteristics of CML patients samples and relative viability of Progenitor Stem Cells and Leukemic Stems Cells upon midostaurin 300 nM exposure.

**Table S8.** Excel file containing FLT3 levels in Non Responder and Responder patients (GSE280476 and GSE14671) and in Leukemic Stems Cells and Leukemic Progenitor Cells (GSE43754).

## Materials and Methods

### Cell culture

K562 cell line was provided by courtesy of professor D. Barilà. LAMA84 cell line was obtained from DSMZ. K562-R cells were generated by exposure of K562 cells to increasingly higher concentration of imatinib during a period of several weeks. The cells were cultured in RPMI 1640 medium (Hyclone, Thermo Scientific, Waltham, MA) supplemented with 10% heat-inactivated fetal bovine serum (ECS0090D Euroclone, Italy, MI), 100 U/ml penicillin and 100 mg/ml streptomycin (Gibco 15140122), 1 mM sodium pyruvate (Sigma-Aldrich, St. Louis, Missouri, United States, S8636) and 10 mM 4-(2-hydroxyethyl)-1-piperazineethanesulfonic acid (HEPES) (Sigma H0887). Responders and non-responder patients derived primary blasts mRNA were provided by courtesy of professor P. Chiusolo. Imatinib (Selleck chemicals, S2475), quizartinib (Selleck chemicals, S1526), ruxolitinib (Selleck chemicals, S1378), MK-2206 (Selleck chemicals, S1078), azacytidine (Selleck chemicals, S1782), venetoclax (Selleck chemicals, S8048), ibrutinib (Selleck chemicals, S2680), copanlisib (Selleck chemicals, S2802) and midostaurin (Selleck chemical, S8064) were used at 0,1-5 µM, 1-100 nM, 0,5-50 µM, 1-100 µM, 1-100 µM, 0,1-50 µM, 10-200 µM, 10-200 nM and 10-500 nM respectively.

### Immunoblot analysis

K562 and K562-R cells were seeded at a concentration of 500.000 cells/ml and treated as indicated. After treatments cells were centrifuged and washed in ice-cold PBS 1x. Next, cells were lysed in ice-cold RIPA lysis buffer (150 mM NaCl, 50 mM Tris–HCl, pH 7.5, 1% Nonidet P-40, 1 mM EGTA, 5 mM MgCl_2_, and 0.1% SDS) supplemented with 1 mM PMSF, 1 mM orthovanadate, 1 mM NaF, protease inhibitor mixture 1x, inhibitor phosphatase mixture II 1x, and inhibitor phosphatase mixture III 1x and incubated for 30 min. Samples were centrifuged at 13,000*g* for 30 min and supernatants were collected. The total protein concentration was determined using the Bradford reagent (Biorad, 5000006). Protein extracts were denatured and heated at 95°C for 10 min in NuPAGE LDS Sample Buffer (Thermo Fisher Scientific, NP0007) and DTT as a reducing agent (NuPAGE Sample Reducing Agent) (Thermo Fisher Scientific, NP0004). Denaturated proteins were resolved using 4–15% Bio-Rad Mini-PROTEAN TGX/CRITERION polyacrylamide gels (Bio-Rad 4561084). Proteins were transferred to Trans-Blot Turbo Mini Nitrocellulose Membranes using a Trans-Blot Turbo Transfer System (Bio-Rad, 17001918). The nitrocellulose membranes were incubated in blocking solution (5% BSA, 0.1% Tween 20 in TBS 1x) at room temperature for 1 hour. Saturated membranes were incubated overnight with primary antibodies diluted in 5% BSA or 5% skimmed milk powder, depending on manufacturer instruction (anti-phospho-c-ABL 1:1000, CST 2865; anti-c-ABL 1:1000, Santa Cruz sc-23; anti-phospho-STAT5 (Y694) 1:1000, CST BK9351; anti-STAT5 1:1000, CST 94205; anti-phospho-ERK1/2 (T202/Y204) 1:1000, CST 9101; anti-ERK1/2 CST 4695; anti-phospho-mTOR (S2448) 1:1000, CST 5536; anti-mTOR 1:1000, CST 4517; anti-phospho-p70 S6 Kinase (T389) 1:1000, CST 9234; anti-p70 S6 Kinase 1:1000, CST 9202; anti-phospho-AKT (S473) 1:1000, CST 4060; anti-AKT 1:1000, CST 9272; anti-GAPDH 1:1000, CST 2118; anti-p62 1:500, Santa Cruz sc-28359; anti-LC3B 1:1000, CST 3868; anti-PI3 Kinase p85α 1:1000, CST 4257; anti-phospho-JAK1 (Y1034/1035) 1:1000, CST 3331; anti-JAK1 1:1000 CST 50996; anti-phospho-FLT3 (Y589/591) 1:1000, CST 3464; anti-FLT3 1:1000, CST 3462; anti-MYC 1:1000, CST 9402; anti-γH2AX 1:5000, CST 9718; anti-NFKB p65 (RELA) 1:500, Santa Cruz sc-8008; anti-ACTIN 1:3000, Sigma A2066; anti-VINCULIN 1:1000, CST 13901; anti-CYCLIN B1 1:1000, CST 4138; anti-CYCLIN H 1:1000, CST 2927; anti-CDK1/2 1:1000, Santa Cruz sc. 53219; anti-CDK2 1:1000, Santa Cruz sc-6248; anti-p21 1:1000, Santa Cruz sc-6246). HRP-conjugated secondary antibodies (Goat Anti-Mouse IgG (H+L)-HRP Conjugate 1:3000, BIORAD 1721011) were diluted in 5% skimmed milk powder, 0.1% Tween 20 in 1× TBS and used for the detection of the primary antibodies. Chemiluminescence was detected using Clarity Western ECL Blotting Substrates (Bio-Rad) and the Chemidoc (Bio-Rad). Band densities were quantified using ImageJ and normalized to the loading control.

### Cell cycle analysis

Cells were treated with imatinib for 24 hours at a concentration of 500.000 cell/ml. The following day 10^6^ cells were collected from each sample and washed with ice-cold PBS. Cells were resuspended in 1 μg/ml DAPI (Thermo Scientific, #62248) and 0.2 mg/ml RNase (Thermo Scientific, #12091021) PBS solution and incubated for 30 minutes before flow cytometry analysis. The fluorescence intensity was detected using CytoFLEX S (Beckman Coulter). Fluorescence intensity was measured using a CytoFLEX S instrument (Beckman Coulter). Weekly quality control checks for the cytometer were performed using CytoFLEX Daily QC Fluorospheres (Beckman Coulter B53230). Data acquisition was conducted using CytExpert software (Beckman Coulter).

### MTT assay

Cell viability was measured using the Cell Proliferation Kit I (MTT) (Roche, 11465007001). Cells were treated as indicated for 24 at a concentration of 50.000 cells/ml or 72 hours at a concentration of 10.000 cells/ml. 100 µl of cell suspension were seeded in technical triplicate in a 96 multiwell plate. After 24 hours of treatment, 10 µl of MTT were added to the cells and incubated for 4 hours at 37 ◦C. Solubilization Buffer was used to dissolve the formazan crystals during an overnight incubation. Finally, the plates were read at 595 nm using a microplate reader (Bio-Rad).

### Real Time quantitative PCR

5·10^6^ cells were centrifuged and resuspended in 1 ml Trizol reagent (Thermo Fisher Scientific). After 5 minutes incubation at room temperature, 200 ul of ice-cold chloroform and vigorously shaken for 15 seconds. Samples were centrifuged at 12000 xg for 15 minutes and aqueous supernatant containing RNA were collected. Next samples were additioned with 500 ul isopropanol and 10 μg of glycogen and incubated overnight at −20°C. The following day samples were centrifuged at 12000 xg for 10 minutes and supernatant was descartes. Pellets were washed with 1 ml ice-cold ethanol 75% and centrifuged again at 7500 xg for 5 minutes. Ethanol was allowed to evaporate and pellets were resuspended in RNase free water. Quantification and purity of the samples was determined with NanoDrop Lite (Thermo Scientific). 1000 ng of RNA from each sample underwent retrotranscription. PrimeScript RT reagent Kit (Takara) was used following manufacturer’s instructions. Specific primers for the BCR::ABL1 gene were designed (forward: 5’-TGACCAACTCGTGTGTGAAACTC-3’; reverse: 5’-TGACCAACTCGTGTGTGAAACTC-3’). RT-qPCR was performed using the SYBR Premix Ex Taq (Takara) kit and the QuantStudio®3 Real-Time PCR instrument (Applied Biosystems). The fold changes in mRNA levels were normalized on actin gene expression. The comparative analysis of gene expression was evaluated by expressing the values as Log_10_2^-ΔCq^.

### Immunofluorescence analysis

To assess γH2AX modulation upon 1µM imatinib 24h treatment, 5·10^6^ cells were treated as indicated, washed in PBS and fixed in 4% PFA for 30 minutes. Next, cells were washed in PBS and permeabilized in a PBS + 0,1% Triton solution for 10 minutes. After the incubation time, cells were centrifuged at 6000g for 5 minutes, washed in PBS and centrifuged again at 10.000 xg for 5 minutes. Cells were blocked in a PBS + 2% BSA + 0,01% Tween20 solution for 1 hour and then centrifuged at 10.000 xg for 5 minutes. Cells were incubated with γH2AX primary antibody (CST 9718, diluted 1:200 in PBS + 2% BSA + 0,01% Tween20) for 1 hour and washed three times in PBS + 0,01% Tween20. Next, cells were incubated with secondary antibody (SouthernBiotech 4050-30, diluted 1:200 in PBS + 2% BSA + 0,01% Tween20) for 1 hour and washed three times in a PBS + 0,01% Tween20 solution. Cells were stained with DAPI (Thermo Scientific #62248, 1:1000 in PBS1X) for 15 minutes, centrifuged at 6000 xg for 5 minutes and mounted for imaging. The experiment was performed in biological triplicate. Samples were quantified considering the percentage of γH2AX-positive cells (> 5 γH2AX foci) over the total number of cells.

### Sample preparation for proteomic and phosphoproteomic analysis

Cells were lysed in SDC lysis buffer additioned with 4% (w/v) SDC, 100 mM Tris-HCl (pH 8.5). Next, samples were boiled at 95° for 5 minutes and sonicated in Bioruptor for 10 cycles at high intensity 30s on/30s off. Protein concentration was determined by BCA assay. inStageTip (iST) method was used for proteome preparation^32^. Briefly, 50µg of protein extract were diluted in 2% SDC buffer and 1% t*rifluoroacetic acid* (*TFA*). SDBRPS tips were washed with i) 100 µl acetonitrile (ACN), ii) 100 µl of 30% methanol and 1% TFA and iii) 150 µl of 0.2 % TFA by centrifuging tips at 1000 xg for 3 minutes. Samples were loaded onto equilibrated columns and spin at 1000 xg for 10 minutes. SDBRPS tips were washed with i) 100 µl of 1% TFA in ethyl acetate, ii) 100 µl of 1% TFA in isopropanol and iii) 0.2% TFA. For protein elution, we used a buffer containing 80% ACN, 5% NH_4_OH in MilliQ water. Samples were centrifuged at 1000 xg for 4 minutes and concentrated by SpeedVac at 45° for ∼45 minutes. Finally, samples were resuspended in 10μl of a buffer additioned with 2% ACN and 0.1% TFA. EasyPhos workflow was used to prepare phosphoproteomics samples as previously described^33^. Briefly, at least 750 μg of protein extract was diluted in 750 μl of ACN and 250 μl of EP enrichment buffer additioned with 36% TFA and 3mM KH_2_PO_4_. Samples were mixed at 2000 xg for 30 s to clear precipitates, then centrifuged at 20.000 xg for 15 minutes and finally transferred in a 2 ml deep-well plate. TiO_2_ beads were used. For each sample, 12:1 (beads: protein) were weighed out and resuspended in EP loading buffer additioned with 80% ACN and 6% (v/v) TFA. Activated TiO_2_ beads were added to each sample and incubated for 5 minutes at 40° C at 2000 rpm. Next, beads were centrifuged at 2000 xg for 1 minute and supernatants (non-phosphosites) were discarded. The beads were resuspended in 500 μl of EP wash buffer, composed of 60% acetonitrile (ACN) and 1% trifluoroacetic acid (TFA) twice and then transferred to a clean tube/plate. Four additional washes with EP wash buffer were carried out mixing at 2000 rpm for 3 seconds. Following the wash steps, the beads were resuspended in 75 μl of EP transfer buffer (80% ACN, 0.5% acetic acid), transferred onto C8 stage tips (double layer), and spun to dryness at 1000 xg for 5 minutes. Phosphopeptides were eluted in 30 μl of EP elution buffer containing 200 μl of NH4OH and 800 μl of 40% ACN into PCR tubes. Immediately afterward, the samples were concentrated in a SpeedVac at 45°C for 20 minutes. Meanwhile, SDBRPS tips (triple layer) were equilibrated using the following steps: i) 100 μl ACN, ii) 100 μl 30% methanol and 1% TFA, and iii) 150 μl 0.2% TFA. After completion of the SpeedVac, SDBRPS loading buffer (1% TFA in isopropanol) was added to the samples. Subsequently, phosphopeptides were loaded onto equilibrated SDBRPS StageTips and washed sequentially with i) 100 μl 1% TFA in ethyl acetate (EtOAc), ii) 100 μl of 1% TFA in isopropanol, and iii) 150 μl of 0.2% TFA. After the wash steps, phosphopeptides were eluted into clean PCR tubes using a buffer containing 60% ACN and 5% NH4OH. Following another SpeedVac step at 45°C for 30 minutes, phosphopeptides were resuspended in 10 μl of a buffer containing 2% ACN and 0.1% TFA.

### Mass spectrometry analysis

The peptides and phosphopeptides underwent desalting using StageTips and were subsequently separated on a reverse-phase column (50 cm, packed in-house with 1.9-mm C18-Reprosil-AQ Pur reversed-phase beads) (Dr. Maisch GmbH). For single-run proteome analysis, separation occurred over 120 minutes, while for phosphoproteome analysis, it extended to 140 minutes. Following elution, the peptides were subjected to electrospray ionization and analyzed via tandem mass spectrometry using an Orbitrap Exploris 480 instrument (Thermo Fisher Scientific). The instrument operated by alternating between a full scan and multiple high-energy collision-induced dissociation (HCD) fragmentation scans, resulting in a total cycle time of up to 1 second.

### Proteome and Phosphoproteome Data processing

Raw files were analyzed using the Spectronaut software. MS/MS spectra were searched against the Homo sapiens UniProtKB FASTA database (September 2014), with an FDR of < 1% at the level of proteins, peptides and modifications. Enzyme specificity was set to trypsin, allowing for cleavage N-terminal to proline and between aspartic acid and proline. The search included cysteine carbamidomethylation as a fixed modification. Variable modifications were set to N-terminal protein acetylation and oxidation of methionine as well as phosphorylation of serine, threonine tyrosine residue (STY) for the phosphoprotemic samples.

### Proteome and Phosphoproteome Bioinformatics Data Analysis

Bioinformatic analysis was conducted within the Perseus software environment^34^, where statistical analysis of both the proteome and phosphoproteome was executed on logarithmized intensities of quantified values across experimental conditions. Normalization of phosphopeptide intensities consisted in subtracting the median intensity of each sample. To identify significantly modulated proteins and phosphopeptides between conditions, a Student t-test with a permutation-based false discovery rate (FDR) cutoff of 0.05 and S0 = 0.1 was employed. Categorical annotation, such as KEGG pathways, was added in Perseus. To address multiple hypothesis testing, a Benjamini-Hochberg FDR threshold of 0.05 was applied.

### Transcription factors’ enrichment analysis from patients RNAseq data

Patients’ transcriptomic data (GSE216837, GSE14671) were transformed in *SignalingProfiler* 2.0 compliant format. We inferred the activity of transcription factors using *run_footprint_based_analysis* function choosing CollecTRI database^35^ as regulon source and regulon minimal size 20.

### Imatinib treated K562 and K562-R vs control network generation with *SignalingProfiler* 2.0

We run the *SignalingProfiler* 2.0 pipeline for K562 cells exposed to imatinib and K562-R cells to generate two networks linking BCR::ABL1 in inactive state (activity = −1) to 9 cancer hallmark phenotypes (Apoptosis, Proliferation, G1/S transition, DNA repair, DNA fragmentation, G1/S transition, Cell cycle block, Cell cycle exit, Autophagy).

#### Protein activity inference

Proteomic and phospho-proteomic data were processed to make them *SignalingProfiler* 2.0 compliant. Kinase activity was inferred by analyzing the modulation of their target phosphosites between sensitive (resistant) cells and control cells using the *run_footprint_based_analysis* with default parameters. Additionally, regulatory phosphosites’ modulation for kinases, transcription factors, and other signaling proteins was considered through the *phosphoscore_computation* function with default parameters. The resulting scores were combined to derive a final activity score. Protein abundance modulation in proteomic data was also considered as a proxy of activity.

For sensitive cells, K562 and LAMA84 multi-omic data were independently exploited to perform *SignalingProfiler* 2.0 protein activity inference step and the two results were merged. We selected 601 proteins that had the same modulation in the two cell lines. We inferred 244 and 258 kinases, 17 and 22 phosphatases, 214 and 261 transcription factors, and 1684 and 2153 other phosphorylated or modulated in abundance signaling proteins, in imatinib exposed K562 and K562-R cells, respectively.

#### Network generation

For both cell lines, a signaling network was constructed using *SignalingProfiler* 2.0 prior knowledge network (PKN) with direct interactions. The PKN was filtered to retain only interactions involving proteins quantified in the (phospho)proteomics data using the *preprocess_PKN* function. A naïve network connecting BCR::ABL1 to inferred signaling proteins was generated using the *two_layer_naive_network* with default parameters. Then, to keep in the naïve network only the interactions coherent with the proteins’ activity, we applied the *SignalingProfiler* 2.0 two-step multi-shot version of vanillaCARNIVAL optimization^36^ with default parameters. The two networks were connected to 9 cancer hallmarks using the *phenoscore_computation* with default parameters. For K562 cells treated with imatinib only interactions from protein to phenotypes coherent with the phenotypic activity were retained using the *optimize_pheno_network* function with default parameters. We generated a network of 200 nodes and 429 edges for K562 cells treated with imatinib and 710 nodes and 1895 edges for K562-R cells. The two networks are available on NDEX (K562: https://www.ndexbio.org/viewer/networks/e6c7826f-4a6d-11ef-a7fd-005056ae23aa, K562-R: https://www.ndexbio.org/viewer/networks/a0c04e02-4a6e-11ef-a7fd-005056ae23aa)

#### BCR::ABL1 dependent and independent functional circuits identification

To obtain BCR::ABL1 dependent functional circuit (**Fig. 1**) the *pheno_to_start_circuit* of *SignalingProfiler* (v. 2) R package was used, selecting BCR::ABL1 as a starting node and maximum path length 7. To obtain BCR::ABL1 independent functional circuit (**Fig. 2**) the same function was used selecting 23 receptors with opposite regulation between K562-R and K562 cells or present only in K562-R network, ‘Proliferation’ and ‘Apoptosis’ phenotypes as end points and maximum path length 6. The BCR::ABL1 independent functional circuit accounted for 438 nodes and 987 edges. The network is available on NDEX at https://www.ndexbio.org/viewer/networks/73523fe5-4a6f-11ef-a7fd-005056ae23aa. For visualization purposes, we selected only paths with maximum path length 5 (**Fig. 2**), obtaining a network of 60 nodes and 111 edges. The generated optimized networks were displayed on Cytoscape using the RCy3 package (v. 2.14.2). The ‘pheno_layout.xml’ XML file provided within the *SignalingProfiler* 2.0 R package was used to set the network style in Cytoscape.

### FDA-drug targets for hematological malignancies prioritization

#### Druggability Score

For druggable targets prioritization we exploited the BCR::ABL1 independent functional circuit. We first removed nodes with incoherent incoming edges (CARNIVAL activity different than 100 or −100) obtaining a network of 424 and 875 edges. For each node, we computed a *topology score* considering: the network degree (i), the number of paths (maximum length = 10) inhibiting apoptosis (ii) and activating proliferation (iii). To not take into account indirect interactions, for each node we excluded paths with length 1, when longer paths were present. Each score was loghartimized and normalized between 0 and 1 and the average was computed (*topology score*). The *topology score* of each node was multiplied with the *CARNIVAL activity score* of the normalized between −1 and 1, obtaining the *Druggability Score.* Proteins with a positive *Druggability Score* are expected to induce more cell death in K562-R cells than in control cells. In contrast, proteins with a negative *Druggability Score* should exhibit the opposite effect. To identify the FDA-approved drug targets for hematological malignancies we manually associated to the drugs of^16^ the Primary Gene Name of the molecular target (*FDA-drug targets catalogue*) and 13 network nodes were extracted.

#### In vitro validation

For *in vitro* validation, we selected 8 network nodes (BCL2, JAK1, BTK, FLT3, PI3KCB and PI3KC3, DNMT1 and DNMT3A) that had a positive *Druggability Score* and that were present in the FDA-drug targets catalogue. As a negative control, we also considered AKT1 node that had a negative *Druggability Score*. Cell viability assay (see MTT assay paragraph) was performed treating K562 and K562-R for 24 hours with Azacytidine (DNMTi), Venetoclax (BCL2i), Ibrutinib (BTKi), Copanlisib (PI3Ki), Ruxolitinib (JAK1i), Midostaurin (FLT3i), Quizartinib (FLT3i) and MK-2206 (AKTi). For each cell line the half maximal inhibitory concentration (IC50) was computed using IC50 calculator web tool (AAT Bioquest) and the opposite of logarithm was computed and compared with the *Druggability Score*.

### Analysis of drug sensitivity of primary CML bone marrow samples

For drug sensitivity testing of primary patient material, bone marrow mononuclear cells (MNCs) from 3 different patients were used. Patient samples were collected in accordance with the Declaration of Helsinki, following informed consent. MNCs were thawed from a biobank and stained for cell sorting using CD45-PE/Cy7 (Biolegend, clone 2D1), CD34-BV510 (Biolegend, clone 581), CD38-BV785 (Biolegend, clone HIT2), and CD26-FITC (Biolegend, clone BA5b). Leukemic progenitor cells (LPCs, CD38 high /CD26 dim) and leukemic stem cells (LSCs, CD38 dim /CD26 high) were sorted (**Fig.S6**). Samples were cultured for 24 hours in the absence or presence of midostaurin 300nM at 37°C, 5% CO 2, and subsequently analyzed using a CytoSMART™ automated cell counter (Corning), including Trypan Blue-based live/dead discrimination.

### CML patients RNA-seq analysis

RNA was extracted from peripheral blasts of 3 imatinib responder and 4 non responder CML patients (see **Table S8** for clinical characteristics). Samples were obtained upon the patients’ informed consent. Briefly, RNA was prepared from PBMCs using the RNeasy Mini Kit (QIAGEN, Germany). The pellets obtained from buffycoat were resuspended in the RLT buffer. We proceed with the addition of equal volume of 70% Ethanol (EtOH) to our already homogenized sample in RTL. It was spinned at ≥8000 x g (≥10,000 rpm) for 15sec. 700 µl of Buffer RW1 were added and the solution was spinned at ≥8000 x g (≥10,000 rpm) for 15sec. It was emptied carefully, and 500 μl of RPE Buffer were added and the solution was spinned ≥8000 x g (≥10,000 rpm) for 2 minutes. The column was emptied again and placed in a new tube and centrifuged at maximum power for 1 minute to dry.

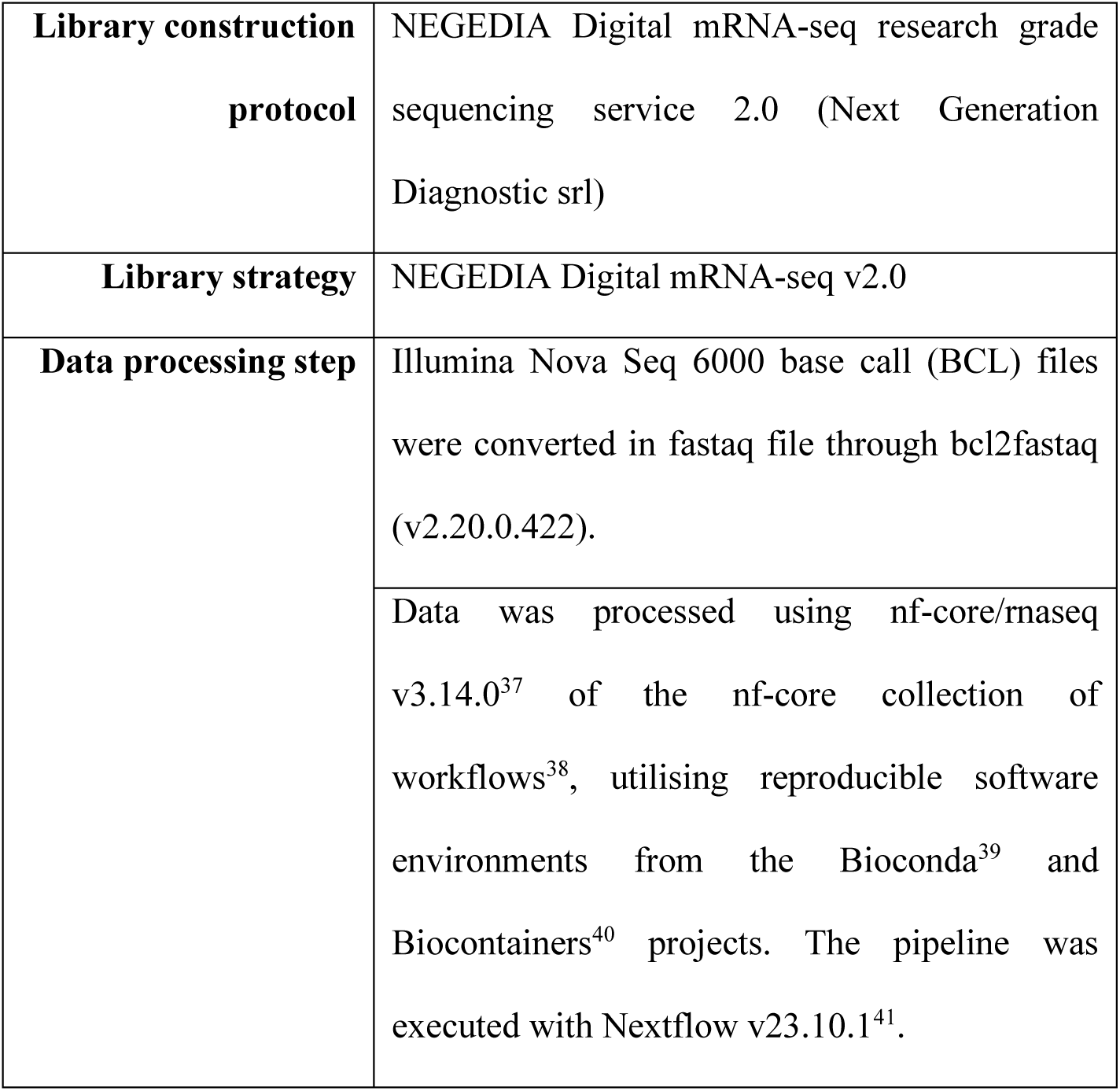

|  |  |
| --- | --- |
| **Library construction protocol** | NEGEDIA Digital mRNA-seq research grade sequencing service 2.0 (Next Generation Diagnostic srl) |
| **Library strategy** | NEGEDIA Digital mRNA-seq v2.0 |
| **Data processing step** | Illumina Nova Seq 6000 base call (BCL) files were converted in fastaq file through bcl2fastaq (v2.20.0.422). |
|  | Data was processed using nf-core/rnaseq v3.14.0<sup>37</sup> of the nf-core collection of workflows<sup>38</sup>, utilising reproducible software environments from the Bioconda<sup>39</sup> and Biocontainers<sup>40</sup> projects. The pipeline was executed with Nextflow v23.10.1<sup>41</sup>. |

The patient transcriptomic data are available on the GEO database under the ID: GSE280476.

### Statistics

All experiments were independently replicated at least three times (n = 3). Data are expressed as means ± standard error (SEM). When comparing three or more groups, statistical analyses were conducted using either one-way or two-way analysis of variance (ANOVA). For comparisons between two groups, the unpaired t-test was employed, assuming a two-tailed distribution. Statistical significance was defined as follows: *p < 0.05; **p < 0.01; ***p < 0.001. Prism 7 (GraphPad) was utilized for all statistical analyses.

