## Supplementary figures S1-S6 for "Phosphoproteomics reveals novel BCR::ABL1-independent mechanisms of resistance in chronic myeloid leukemia"

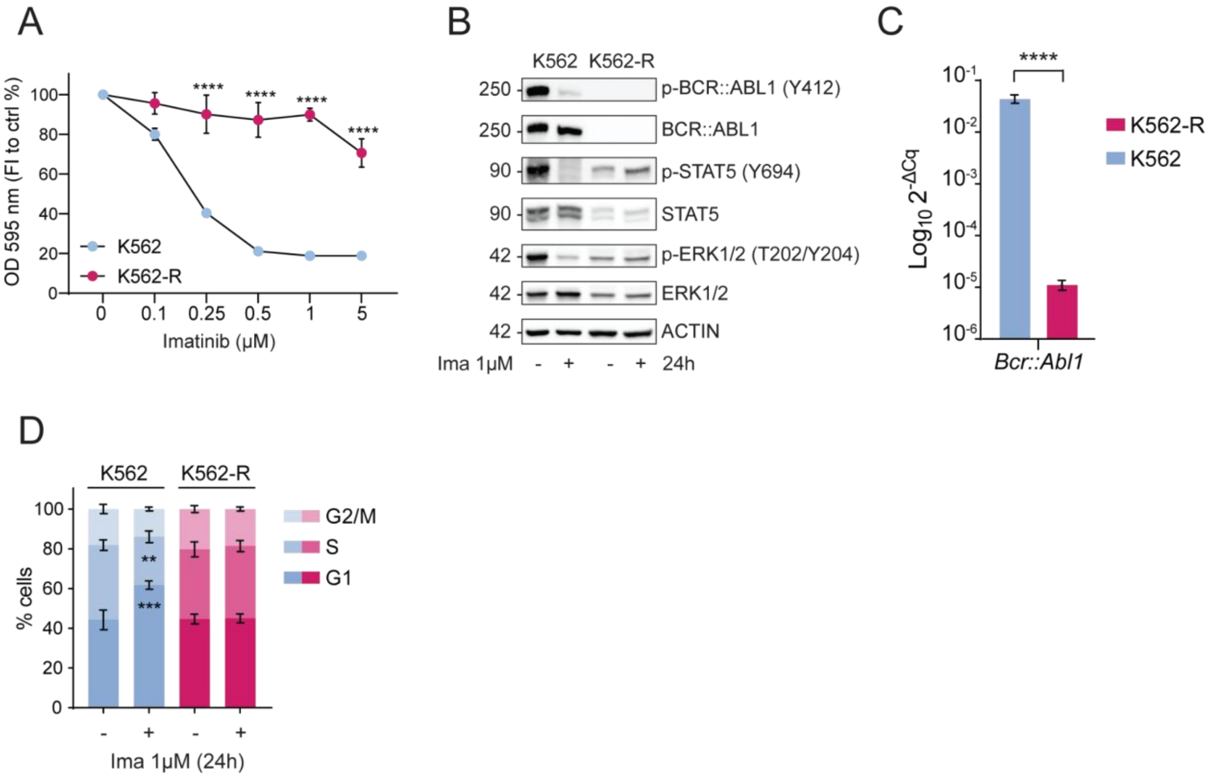


**Figure S1. Characterization of K562-R cell line**

**A.** MTT viability assay. Sensitive cells (blue) and resistant cells (red) were exposed to increasing concentration of imatinib (0.1µM, 0.25µM, 0.5µM, 1µM and 5µM) for 72 hours. The graph shows the percentage of absorbance at 595nm normalized on control condition.

**B.** Representative western blots of BCR::ABL1 activity status (Y412), MAPK and JAK/STAT canonical downstream pathways (p-STAT5 (Y694), p-ERK1/2 (T202/Y204)) in K562 and K562-R cells upon 24h of 1µM imatinib treatment.

**C.** Real Time quantitative PCR was performed to measure BCR::ABL1 transcript levels in K562 and K562-R cells. Bar graph shows quantification of the BCR::ABL1 mRNA levels as $\mathrm{Log10}_2{}^{-\Delta Cq}$.

**D.** Flow cytometry analysis of cell cycle progression of K562 and K562-R cells upon 24h exposure to imatinib 1 µM. The bar graph shows the percentage of cells in a specific cell cycle phase measured using DAPI staining.

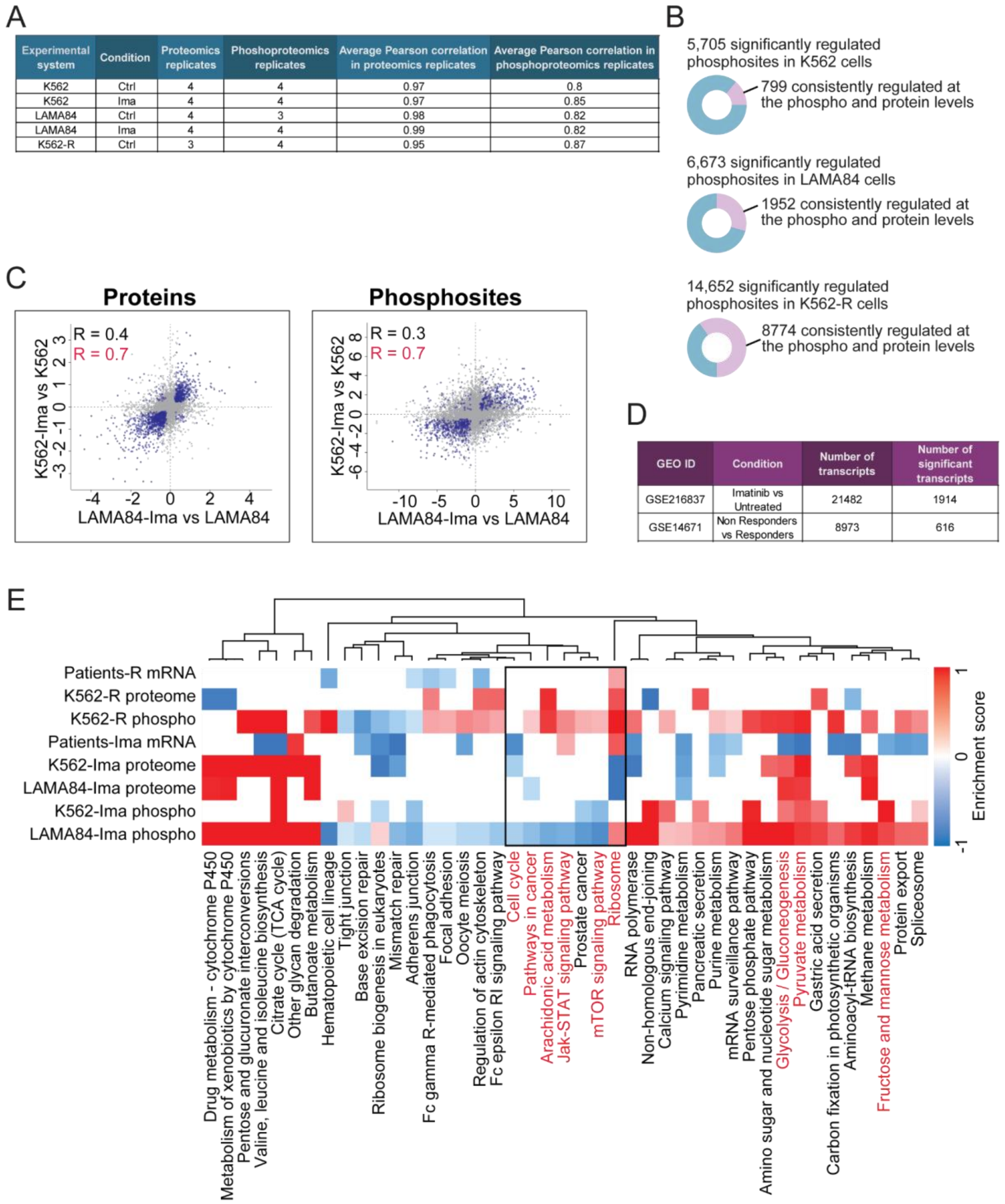


**Figure S2. Technical and functional overview of CML multi-omics datasets**

**A.** Table reporting the number of replicates and the average Pearson correlation for K562, LAMA84 and K562-R cells in control condition and upon imatinib exposure in proteomics and phosphoproteomics experiments.

**B.** Pie charts showing the proportion of phosphopeptides consistently modulated by imatinib (upper panels) or in K562-R cells at their phosphorylation and protein levels.

**C.** Scatterplots showing the Pearson correlation coefficients between the imatinib-dependent changes at the phosphoproteome and proteome levels of K562 cells as compared to LAMA84 cells. R indicates Pearson correlation.

**D.** Table reporting the GEO ID of datasets employed for independent validations. For each dataset is reported the total and the significant number of quantified transcripts.

**E.** Heatmap representing the enrichment score of KEGG pathways significantly up-regulated (red) or down-regulated (blue) by imatinib treatment at the phosphoproteome, proteome and transcriptome levels in sensitive and resistant CML cells.

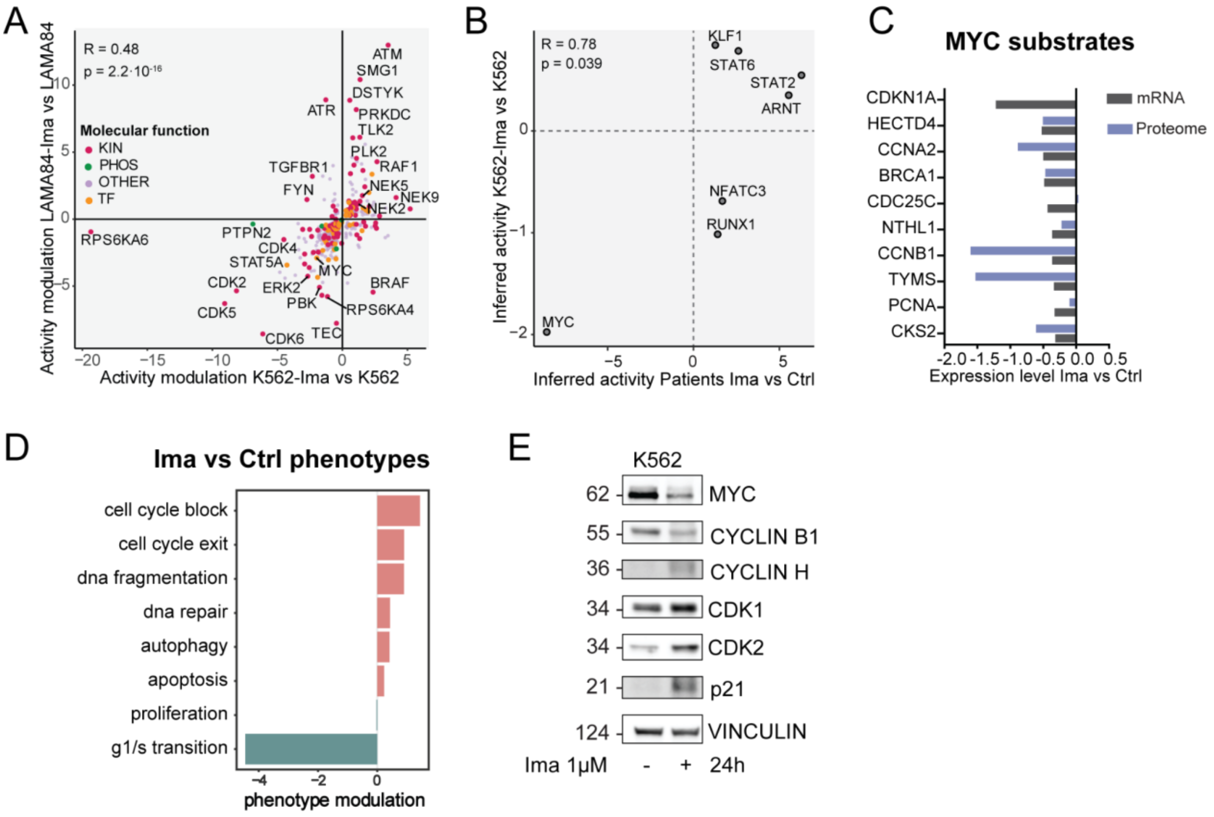


**Figure S3. Functional analysis of imatinib-treated sensitive cell models and patients**

**A.** Scatterplot showing the comparison between imatinib-mediated changes of inferred protein activity in K562 (x-axis) and LAMA84 (y-axis) cells. Each dot represents a protein, and the color indicates the molecular function: kinases (red), phosphatases (green), others (violet), and transcription factors (orange). R indicates Pearson correlation

**B.** Scatterplot of transcription factors' inferred activity with *SignalingProfiler* 2.0 in imatinib treated patients-derived blasts and K562 cell line (y-axis). R indicates Pearson correlation.

**C.** Bar graph reporting K562 protein (blue) and patients mRNA (gray) abundance of MYC targets upon imatinib exposure.

**D.** Bar plot of the phenotypic modulation upon imatinib treatment in sensitive cells inferred by *SignalingProfiler 2.0*. Blue and red bars represent inactive and active phenotypes, respectively.

**E.** Representative western blot of how proteins involved in cell cycle regulation are modulated by 24h imatinib treatment.

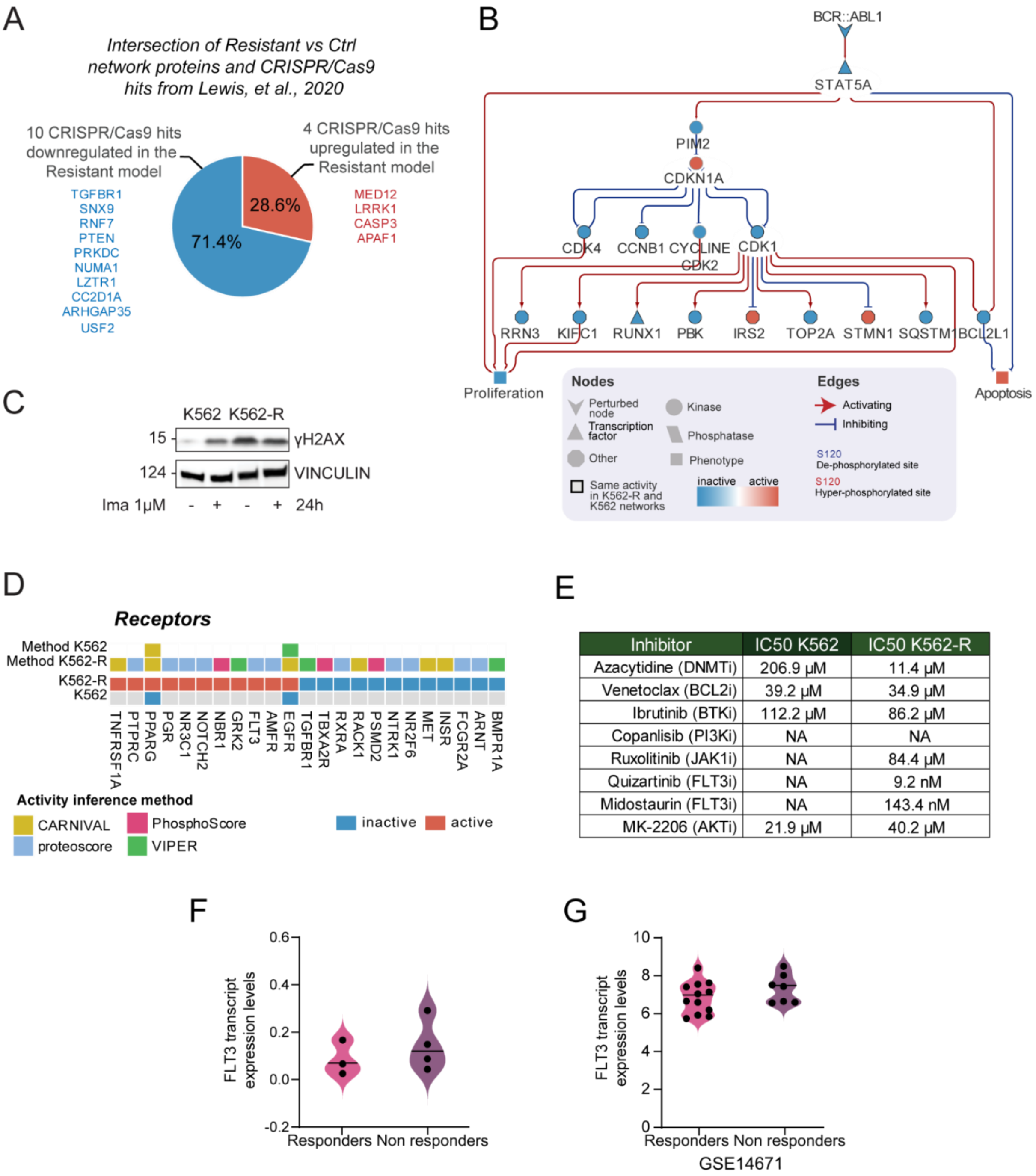


| Inhibitor | IC50 K562 | IC50 K562-R |
|---|---|---|
| Azacytidine (DNMTi) | 206.9 μM | 11.4 μM |
| Venetoclax (BCL2i) | 39.2 μM | 34.9 μM |
| Ibrutinib (BTKi) | 112.2 μM | 86.2 μM |
| Copanlisib (PI3Ki) | NA | NA |
| Ruxolitinib (JAK1i) | NA | 84.4 μM |
| Quizartinib (FLT3i) | NA | 9.2 nM |
| Midostaurin (FLT3i) | NA | 143.4 nM |
| MK-2206 (AKTi) | 21.9 μM | 40.2 μM |

**Figure S4. Construction and validation of K562-R resistance model**

**A.** Pie chart of the intersection of K562-R cells signaling network proteins with CRISPR/Cas9 hits from Lewis, et al., 2020 work.

**B.** BCR::ABL1-dependent mechanisms in K562-R cells signaling network. Functional submodel extracted from SignalingProfiler 2.0 output linking BCR::ABL1 to proteins with the same modulation in K562-R and K562 upon imatinib treatment (highlighted in black).

**C.** Representative western blot showing protein levels modulation of DNA damage marker γH2AX in control cells and K562-R cells upon 24h imatinib perturbation.

**D.** Heatmap reporting for control and K562-R cells the activity of receptors in SignalingProfiler 2.0 generated networks.

**E.** Table showing IC50 calculations derived from MTT viability assays of FDA-approved inhibitors on K562 and K562-R for 24 hours.

**F.** Quantification of FLT3 transcript levels in RNAseq analysis of responder and non responder patients derived primary blasts (GSE280476).

**G.** Quantification of FLT3 transcript levels in responders and non responders CML patients from RNAseq analysis obtained from GEO dataset (GSE14671).

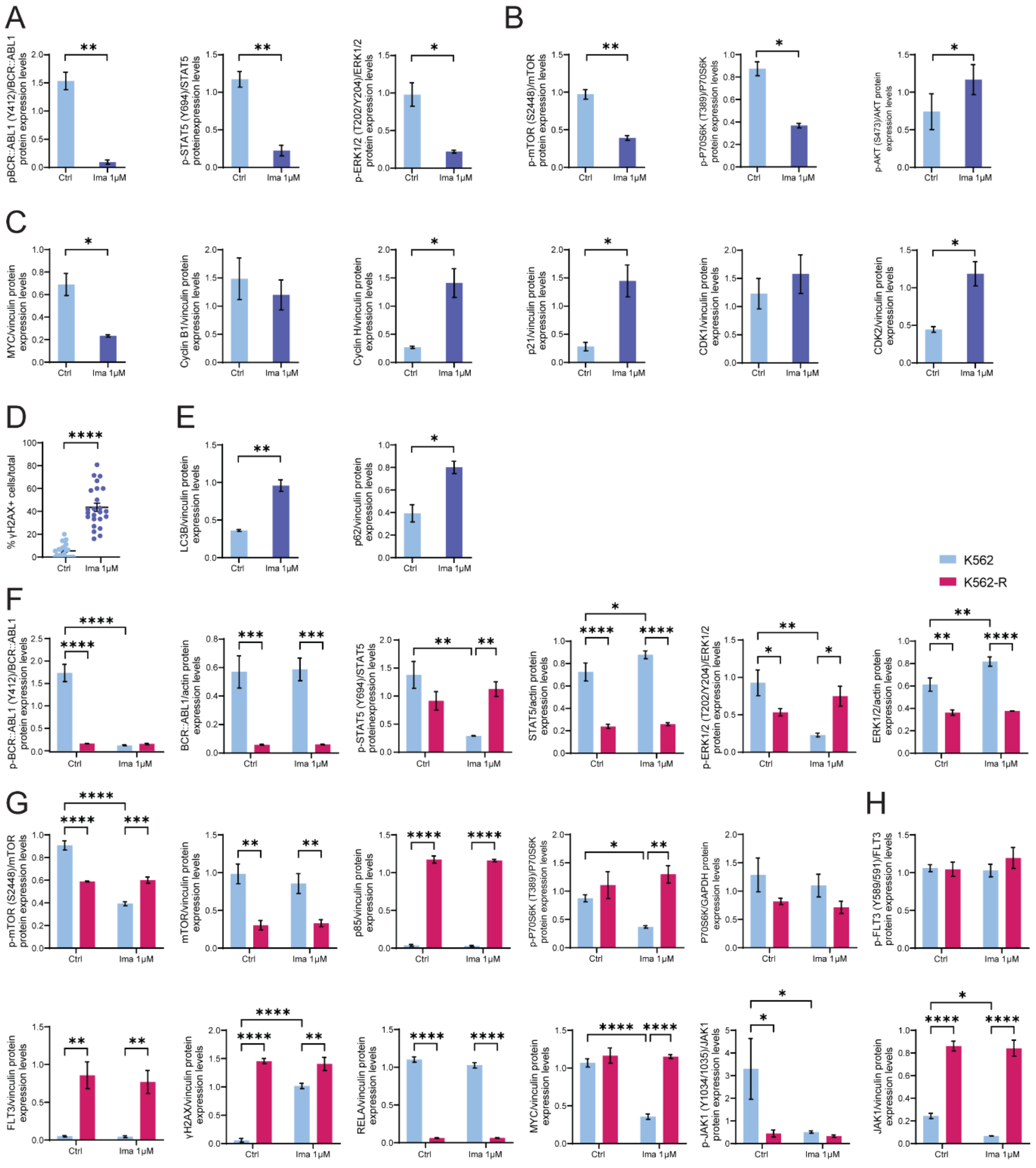


**Figure S5. Quantification of proteins phosphorylation status and expression levels**

**A.** Quantification of phosphorylation status of p-BCR::ABL1 (Y412), p-STAT5 (Y694), p-ERK1/2 (T202/Y204) upon 24h of 1µM imatinib treatment in K562 cells.

**B.** Quantification of phosphorylation status of p-mTOR (S2448), p-P70S6K (T389) and p-AKT (S473) upon 24h of 1µM imatinib treatment in K562 cells.

**C.** Quantification of protein abundance of MYC, Cyclin B1, Cyclin H, p21, CDK1 and CDK2 upon 24h of 1µM imatinib treatment in K562 cells.

**D.** Quantification of the percentage of γH2AX+ K562 cells upon 24h exposure to imatinib. Cells were considered positive if they show ≥ 5 foci. The number of γH2AX positive cells were normalized on the total number of cells.

**E.** Quantification of protein abundance of key autophagy regulators LC3B and p62 in K562 cells upon 90 minutes imatinib treatment.

**F.** Quantification of phosphorylation status and protein abundance of p-BCR::ABL1 (Y412), p-STAT5 (Y694), p-ERK1/2 (T202/Y204) upon 24h of 1µM imatinib treatment in K562 and K562-R cells.

**G.** Quantification of phosphorylation status and protein abundance of p-mTOR (S2448), p85α and p-P70S6K (T389) upon 24h of 1µM imatinib treatment in K562 and K562-R cells.

**H.** Quantification of phosphorylation status and protein abundance of p-FLT3 (Y589/591), γH2AX, RELA, MYC and p-JAK1 (Y1034/1035) upon 24h of 1µM imatinib treatment in K562 and K562-R cells.

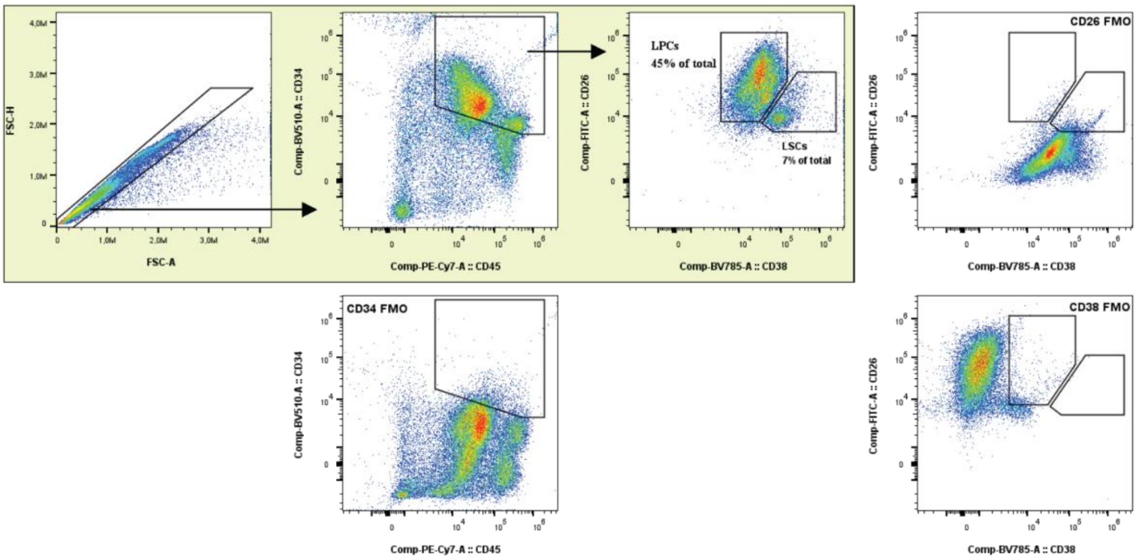


**Figure S6. Gating strategy for sorting CD38+ cells (LPCs) and CD26+ cells (LSCs) from patients derived samples**
